# A model of preferential pairing between epithelial and dendritic cells in thymic antigen transfer

**DOI:** 10.1101/2021.09.13.460045

**Authors:** Matouš Vobořil, Jiří Březina, Tomáš Brabec, Jan Dobeš, Ondřej Ballek, Martina Dobešová, Jasper Manning, Richard S. Blumberg, Dominik Filipp

## Abstract

Medullary thymic epithelial cells (mTECs) which produce and present self-antigens are essential for the establishment of central tolerance. Since mTEC numbers are limited, their function is complemented by thymic dendritic cells (DCs), which transfer mTEC-produced self-antigens via cooperative antigen transfer (CAT). While CAT is required for effective T cell selection, many aspects remain enigmatic. Given the recently described heterogeneity of mTECs and DCs, it is unclear whether the antigen acquisition from a particular TEC subset is mediated by preferential pairing with specific subset of DCs. Using several relevant Cre-based mouse models controlling the expression of fluorescent proteins, we found that in regards to CAT, each subset of thymic DCs preferentially targets distinct mTEC subset(s) and importantly, XCR1^+^ activated DCs represented the most potent subset in CAT. Interestingly, one thymic DC can acquire antigen repetitively and of these, monocyte-derived DCs (moDC) were determined to be the most efficient in repetitive CAT. moDCs also represented the most potent DC subset in the acquisition of antigen from other DCs. These findings suggest a preferential pairing model for the distribution of mTEC-derived antigens among distinct populations of thymic DCs.

## Introduction

Central tolerance, which operates during T-cell development in the thymus, can result in the elimination of self-reactive T-cells or their deviation into thymic regulatory T-cell (tTreg) lineage (Klein et al., 2019). The underlying principle of this event compels immature T-cells to test their T-cell receptor (TCR) for potential self-reactivity through scanning of self-antigens which are presented by antigen presenting cells (APCs). Among all thymic APCs, thymic epithelial cells (TECs) are central in this selection process (Klein et al., 2014). Based on their localization within the thymus, TECs are generally divided into two major populations: cortical TECs (cTEC) and medullary TECs (mTECs) (Derbinski et al., 2001). Recently, single-cell RNA sequencing (scRNAseq) revealed an unexpected heterogeneity of mTECs with at least five distinct subsets defined by their developmental stage, transcription profile, and function (referred to as mTEC-I, -II, -IIIa, IIIb, and Tuft cells) (Baran-Gale et al., 2020; Bornstein et al., 2018; Miller et al., 2018).

Due to their unique ability to express and present more than 80% of the protein-coding genome, mTECs are well-adapted to serve as a principal self-antigen-producing cellular component of central tolerance (Brennecke et al., 2015; Meredith et al., 2015; Sansom et al., 2014). This is facilitated, in part, by the expression of the Autoimmune regulator (Aire). Aire controls the gene expression of a large set of tissue restricted antigens (TRAs) found only in the immune periphery (Derbinski et al., 2001). Interestingly, an effective display of a complete set of thymically expressed TRAs is achieved by their combinatorial mosaic expression by each mTEC with any particular TRA expressed by only 1-3% of mTECs (Derbinski et al., 2008) while a single mTEC is capable of expressing up to 300 different TRAs (Meredith et al., 2015; Sansom et al., 2014). However, mTEC subsets are not equal in terms of Aire expression and TRA presentation. During their progression through mTEC-I, -II, -IIIa, and -IIIb stages, the highest Aire and TRA expression is observed in mTEC-II, historically referred to as mTEC^high^. As mTEC-II enter pre-post Aire and post-Aire phases (phase -IIIa and -IIIb, respectively), they downregulate the expression of Aire, although their TRA protein levels remain high, making them available for further use by other cells (Kadouri et al., 2020). The extent of the expression of TRA in mTEC-I (referred to as mTEC^low^) is limited (Baran-Gale et al., 2020; Bornstein et al., 2018).

The relatively low number of mTECs in comparison to the sheer number of developing T-cells, coupled to mosaic and stage-restricted expression of TRAs, places significant constraints on the process of T cell selection. To overcome this limitation, TRAs from apoptotic mTECs can be transferred into, and indirectly presented to T-cells, by thymic dendritic cells (DCs) via the process of cooperative antigen transfer (CAT) (Gallegos & Bevan, 2004; Koble & Kyewski, 2009). It has been demonstrated that CAT is critical for the establishment of central tolerance to mTEC-derived self-antigens (Lancaster et al., 2019; Perry et al., 2014; Perry et al., 2018). Despite its importance, the elucidation of its basic principles has been hampered by the complexity of thymic DC populations.

In general, thymic DCs can be divided into two major categories – plasmacytoid DCs (pDCs) and classical DCs (cDCs), the latter of which can be subdivided into cDC1 and cDC2 subsets (Guilliams et al., 2014). Previous studies have shown that these DC subsets vary in their capacity to acquire mTEC-derived antigens (Kroger et al., 2017; Lancaster et al., 2019; Vobořil et al., 2020). cDC1s were shown to strongly acquire GFP antigen from mTEC in *Aire-GFP* mouse model (Perry et al., 2018). On the other hand, the cDC2 subset robustly acquires mOVA antigen in the *RIP-mOVA* mouse model (Lancaster et al., 2019). Since the expression of Aire-driven GFP and mOVA in the thymus was largely restricted to Aire^+^ mTECs (Gardner et al., 2008) and mTEC^Low^/post-Aire mTECs, respectively, (Mouri et al., 2017), it has been inferred that distinct subsets of thymic DCs acquire antigens from distinct subsets of mTECs. However, our recent scRNAseq analysis along with data from the human thymus cell atlas study unearthed a much broader heterogeneity of DCs in the thymus of mice and humans (Park et al., 2020; Vobořil et al., 2020). Thus, a more comprehensive analysis is needed to determine the mode of CAT between defined subsets of TECs and DCs as well as other means of thymic antigen spreading.

In this study, we used several Cre reporter mouse models in which the expression of fluorescent TdTOMATO protein (TdTOM) is restricted to different subsets of TECs. We present evidence suggesting that distinct subsets of thymic DCs preferentially acquire TdTOM from a certain subset of TECs. Using the Confetti^Brainbow2.1^ system, we have also shown that CAT can occur as a repetitive event whereby a single thymic CD11c^+^ cell can acquire antigen from two or more individual TECs. Furthermore, based on our data, we postulate that antigen transfer can also occur between DC subsets themselves. Thus, this dataset suggests a deterministic model of preferential engagement of specific mTEC and DC subsets for directional thymic antigen spreading.

## Results

### Thymic epithelial cell models of cooperative antigen transfer

The robustness of scRNAseq has yielded in recent years comprehensive knowledge in regards to detailing thymic APCs inventory as well as a list of suitable markers (Baran-Gale et al., 2020; Bautista et al., 2021; Bornstein et al., 2018; Dhalla et al., 2020; Park et al., 2020; Vobořil et al., 2020; Wells et al., 2020). The combinatorial specificity of these markers has led us to design novel flow cytometry gating strategies that allow us to study CAT.

To understand antigen transfer trajectories within the intricate network of all subsets of TECs and CD11c^+^ APCs identified thus far, we first established mouse models where cytoplasmic expression of TdTOM is preferentially confined to distinct TEC subsets. By crossing three previously characterized Cre-based mouse models with a ROSA26^TdTOM^ mouse strain, we generated: (i) Foxn1^Cre^ROSA^26TdTOM^ (Foxn1^Cre^) mice which express TdTOM in all populations of CD45^−^EpCAM^+^ TECs (Gordon et al., 2007; Vobořil et al., 2020), (ii) Csnβ^Cre^ROSA26^TdTOM^ (Csnβ^Cre^) with Casein β (Csnβ) loci operating as an Aire-independent TRA which confines TdTOM expression to mTEC^High^ subset and their closest progenitors and progeny (Bornstein et al., 2018; Tykocinski et al., 2010), and (iii) Defa6^iCre^ROSA26^TdTOM^ (Defa6^iCre^). The latter model represents the “classical” Aire-dependent TRA model, in which TdTOM is expressed in 1-3% of Aire^+^ mTEC^High^ cells as well as Post-Aire mTEC progeny (Adolph et al., 2013; Dobeš et al., 2015) (Figure 1b-c).

**Figure 1.**
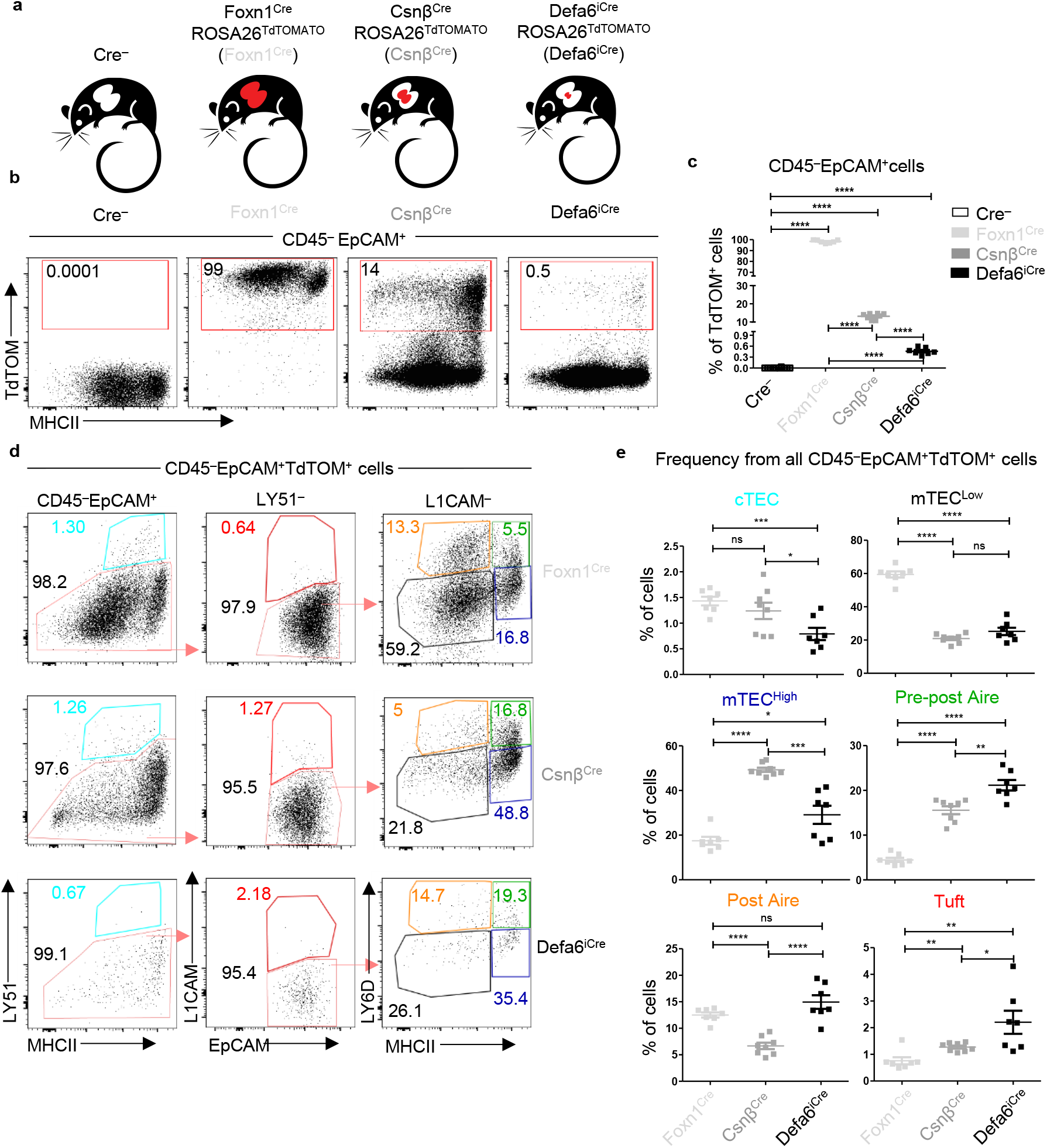
The phenotype and frequency of TEC subsets in Cre-based mouse models of CAT. **(a)** Mouse models of CAT with confined expression of TdTOM to distinct TEC subsets. **(b)** Representative flow cytometry plots showing the frequency of TdTOM^+^ cells among CD45^−^EpCAM^+^ cells isolated from a MACS-enriched CD45^−^ thymic population from Foxn1^Cre^ROSA26^TdTOM^ (Foxn1^Cre^), Csnβ^Cre^ROSA26^TdTOM^ (Csnβ^Cre^) and Defa6^iCre^ROSA26^TdTOM^ (Defa6^iCre^) mice. **(c)** Quantification of TdTOM^+^ cells from Fig. 1b (mean ± SEM, *n*=7-12 mice from 3 independent experiments). **(d)** Representative comparative flow cytometry plots of different TEC subsets in Foxn1^Cre^, Csnβ^Cre^ and Defa6^iCre^ mice. **(e)** Quantification of TEC subset frequencies from plots in Fig. 1d (mean ± SEM, *n*=7-8 mice from 3 independent experiments). Statistical analysis in (c) and (e) was performed using an unpaired, two-tailed Student’s t-test, p≤0.05 = *, p≤0.01 = **, p≤0.001***, p<0.0001 = ****, ns = not significant.

The gating strategy implemented to assess the frequency of TdTOM-labelled CD45^−^EpCAM^+^ TEC subsets in the Cre models introduced above (Figure 1a-c) is shown in Supplementary Figure 1a. Six subsets of TECs were distinguished: cTEC, mTEC^Low^, mTEC^High^, two subsets of LY6D^+^ terminally differentiated subsets: Pre-post Aire and Post-Aire mTECs, and L1CAM^+^ thymic Tuft cells. The results confirmed the differences among the TEC subsets found within the thymic CD45^−^EpCAM^+^TdTOM^+^ population in the three generated mouse models. Whereas cTEC and mTEC^Low^ subsets were overrepresented in Foxn1^Cre^, and the mTEC^High^ subset in the Csnβ^Cre^ model, the frequencies of Pre-post Aire, Post-Aire, and Tuft mTECs were increased in the Defa6^iCre^ model (Figure 1d-e). This data validated the utility of the Cre-based ROSA26^TdTOM^ mouse models to study CAT, since the expression of TdTOM protein was in each model predictably enriched in different subsets of TECs.

### Antigen transfer of TdTOM to thymic dendritic cells

Having characterized the distinct distribution of TdTOM in TEC subsets in our Cre-based ROSA26^TdTOM^ mouse models, we next tested the distribution of the TdTOM among its acceptors, the thymic population of CD11c^+^ cells (Figure 2a). As shown previously (Vobořil et al., 2020) and in Figures 2b and c, TdTOM is mostly acquired by CD11c^+^ cells. The robustness of this transfer which is heavily dependent on the type of Cre-based ROSA26^TdTOM^ mouse model was then examined. Whereas TdTOM positivity was observed in ∼6% of CD11c^+^ cells in the Foxn1^Cre^ model, its frequency in Csnβ^Cre^ and Defa6^iCre^ was limited to ∼0,6% and ∼0,02%, respectively (Figure 2c). Interestingly, even though the frequency of TdTOM^+^ TECs was significantly decreased across the Foxn1^Cre^, Csnβ^Cre^ and Defa6^iCre^ mouse models (Figure 1c), the ratio between the frequency of TdTOM^+^CD11c^+^ and TdTOM^+^CD45^−^EpCAM^+^ TECs cells in the models used was comparable (Figure 2d). These analyses argue for the similarity in CAT efficiency between donor TECs and CD11c^+^ APC acceptors, irrespective of the robustness and cell-subset range of TdTOM expression in TECs.

**Figure 2.**
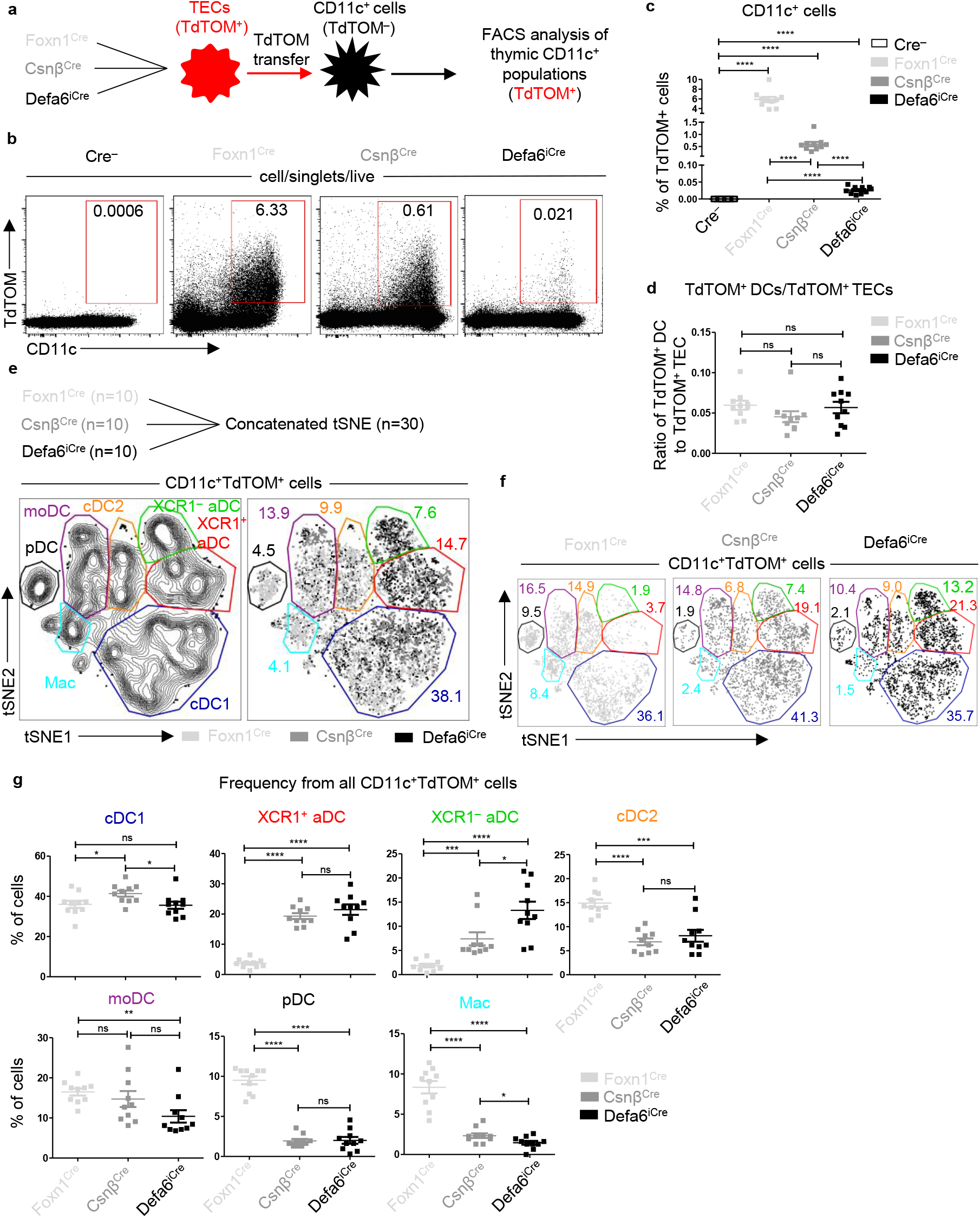
Antigen transfer of TdTOM to thymic dendritic cells. **(a)** Experimental design. **(b)** Representative flow cytometry plots comparing the frequency of TdTOM^+^CD11c^+^ cells among MACS-enriched CD11c^+^ thymic cells from mice models described in (a). **(c)** Quantification of TdTOM^+^CD11c^+^ cells from (b) (mean ± SEM, *n*=10 mice from minimum of 3 independent experiments). **(d)** Comparison of the ratio between the frequency of TdTOM^+^CD11c^+^ (quantified in c) to TdTOM^+^ TEC (quantified in Fig. 1c) subsets in mice models described in (a) (mean ± SEM, *n*=10 mice from minimum of 3 independent experiments). **(e)** Concatenated (n=30 mice) and **(f)** separate (n=10 mice) flow cytometry tSNE analysis of TdTOM^+^CD11c^+^ cells from the three mice models described in (a). **(g)** Quantification of TdTOM^+^CD11c^+^ subset frequencies described in (e) (mean ± SEM, *n*=10 mice from minimum of 3 independent experiments). Statistical analysis in (c), (d), and (g) was performed using an unpaired, two-tailed Student’s t-test, p≤0.05 = *, p≤0.01 = **, p≤0.001***, p<0.0001 = ****, ns = not significant.

To study CAT in the mouse models defined above, we determined seven subpopulations of thymic CD11c^+^ cells (Supplementary Figure 2a). These cells are comprised of three major categories: B220^+^ plasmacytoid DCs (pDC), CD11c^Low^MHCII^Low^CX3CR1^+^ macrophage-like population (Mac), and CD11c^+^MHCII^High^ cells which represent a conventional type of thymic DCs. Historically, the thymic DCs were subdivided into two groups, cDC1 and cDC2, defined by the expression of chemokine receptor, XCR1 and SIRPα, respectively (Li et al., 2009). Recently, the SIRPα^+^ DCs were described to encompass a minimum of two different subpopulations, defined by the expression of MGL2 (CD301b) and CD14 to MGL2^+^CD14^−^ cDC2 and MGL2^+^CD14^+^ monocyte-derived DCs (moDC) (Vobořil et al., 2020). It has also become evident that DCs could be phenotypically and functionally defined by their activation status (Ardouin et al., 2016; Park et al., 2020; Vobořil et al., 2020). Hence, two phenotypically distinct subsets of activated DC (aDCs), CCR7^+^XCR1^+^ and CCR7^+^XCR1^−^, can be identified (Supplementary Figure 2a). A comparative analysis of the capacity of each of these thymic CD11c^+^ APC subsets to acquire TEC-derived TdTOM showed that consistent with previously published data (Ardouin et al., 2016), XCR1^+^ aDCs were the most efficient cells involved in CAT irrespective of the Cre-based ROSA26^TdTOM^ model used. On the other hand, while Macs and pDCs were relatively inefficient, the remaining subsets varied in this efficiency depending on the Cre-model analyzed (Supplementary Figure 2b). Using bone marrow (BM) chimeras of sub-lethally irradiated mouse models (Foxn1^Cre^, Csnβ^Cre^ and Defa6^iCre^) reconstituted with congenically marked BM cells isolated from WT animals, we verified that TdTOM is indeed transferred from TECs to all subpopulations of thymic CD11c^+^ APCs and is not endogenously expressed by these APCs themselves (Supplementary Figure 2c-f).

Since the frequency of each CD11c^+^ APC subset as well as their capacity to acquire TEC-derived antigen differ, we next assessed their contribution to CAT in all three Cre-based ROSA26^TdTOM^ mouse models. Due to the comparative nature of this approach (comparing the efficiency of CAT for each CD11c^+^ APC subset in each Cre model), we first performed an unsupervised flow cytometry analysis of all CD11c^+^TdTOM^+^ cells concatenated from 10 independent samples from each of the Cre-based mouse models (30 samples) (Figure 2e). Based on the markers shown in Supplementary Figure 3a, we identified all phenotypically distinguished CD11c^+^ APC subsets in the resulting tSNE plot (Supplementary Figure 3a-b). Analyzing each of the Cre-based ROSA26^TdTOM^ mouse models individually (Figure 2f), the data revealed that whereas the contribution of cDC1s and moDCs to CAT is robust in all the cases studied, the contribution of pDCs, Macs, cDC2s, and both populations of aDC subsets varied among the models. Notably, cDC2s, pDCs, and Macs were significantly increased in the Foxn1^Cre^ mouse model. In contrast, the frequency of XCR1^+^ and XCR1^−^ aDCs was the lowest in Foxn1^Cre^, with an increase detected in Csnβ^Cre^, and the highest frequency detected in Defa6^iCre^ model (Fig. 2g). Taken together, this data shows that the extent of the involvement of each DC subset in CAT depends on the distribution of TdTOM protein expression among the different subtypes of TECs, and/or the overall proportion of TECs expressing the TdTOM. In this way, the assorted expression of TdTOM antigen by a limited but defined subset of TECs allows the visual identification of those DC subsets which engage these TEC subsets during CAT.

### Projecting preferential trajectories of CAT between TEC and thymic DC subsets

To reveal the possible combinations of TEC and DC subsets that are preferentially engaged in CAT, the frequency of TdTOM^+^ TEC subsets shown in Figure 1d-e and TdTOM^+^ thymic DC subsets from Figure 2e-g were visualized as color-coded pie charts for each Cre-based mouse model used (Figure 3a). Upon inspection of these charts, a trend towards the decrease of mTEC^Low^ versus the increase of mTEC^High^ and Pre-post Aire cells from Foxn1^Cre^ to Csnβ^Cre^ to Defa6^iCre^ mouse models is apparent. On the other hand, a decrease in the frequency of pDCs and Macs was observed while the contribution of XCR1^+^ and XCR1^−^ aDCs in the TdTOM^+^ gate was increased. The simplest interpretation of these correlations is the possibility of pDCs and Macs preferentially acquire antigen from mTEC^Low^ subset, while the CAT to both populations of aDCs, is likely associated with mTEC^High^ and Pre-post Aire cells (Figure 3a).

**Figure 3.**
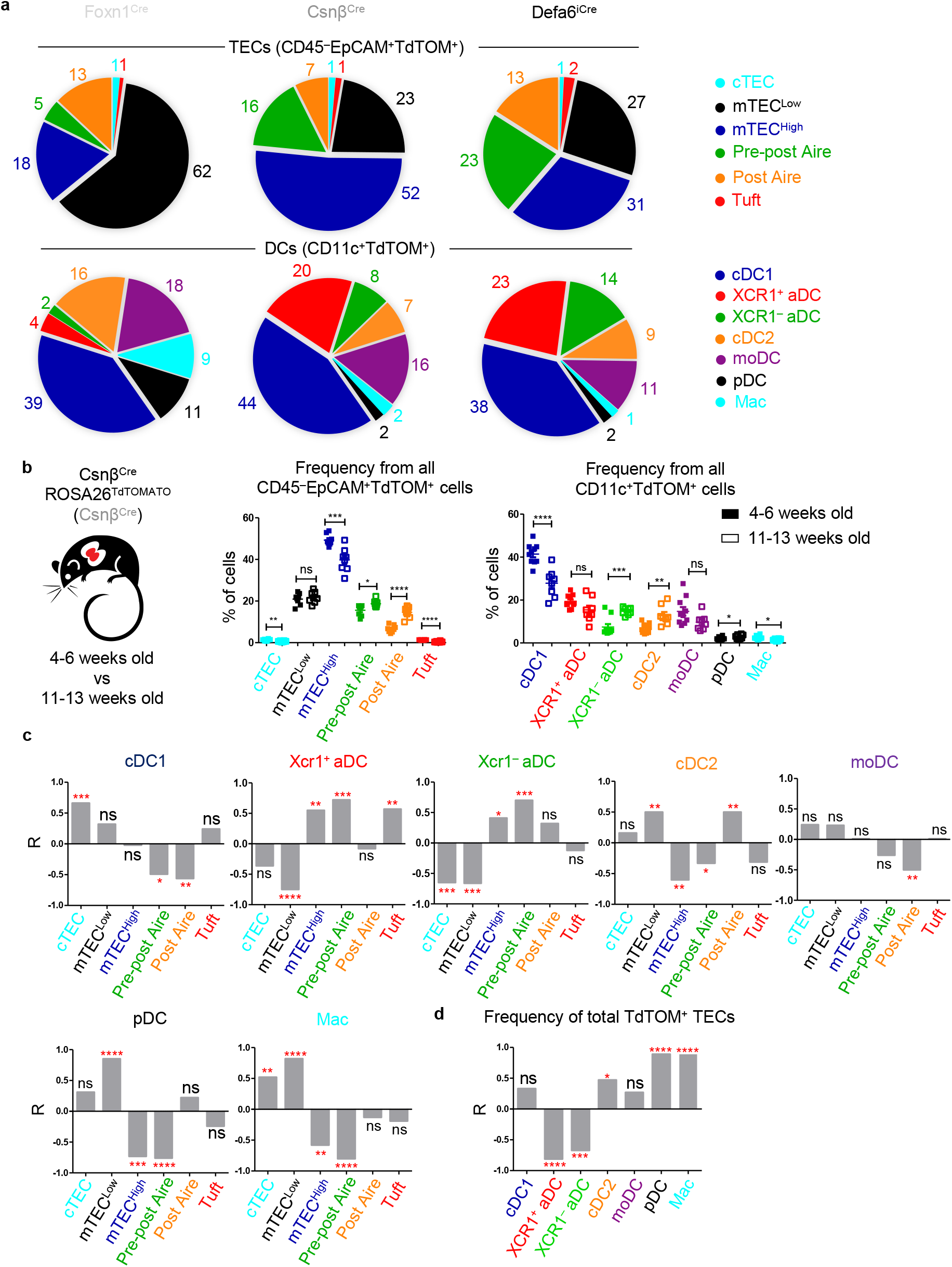
TdTOM antigen transfer to distinct thymic DC subsets correlates with its confined expression in phenotypically defined subsets of TECs. **(a)** Pie chart visualization of the frequency of TEC subsets from all CD45^−^EpCAM^+^TdTOM^+^ cells (from Figure 1e) (upper part) and DC subsets from all CD11c^+^TdTOM^+^ cells (from Fig. 2g) (lower part) from all described mice models. **(b)** Comparison of the frequency of TEC and DC subsets from all TdTOM^+^ cells between young (4-6 weeks old) and older (11-13 weeks old) Csnβ^Cre^ROSA26^TdTOM^ (Csnβ^Cre^) mice (mean ± SEM, *n*=8-10 mice from a minimum of 3 independent experiments). Statistical analysis was performed using unpaired, two-tailed Student’s t-test, p≤0.05 = *, p≤0.01 = **, p≤0.001***, p<0.0001 = ****, ns = not significant. **(c)** Bar charts showing linear regression (R) between the frequencies of TdTOM^+^ TECs and the indicated subset of TdTOM^+^ DCs from (a) and (b) (n=5-8 mice, from a minimum of 3 independent experiments). **(d)** Bar chart showing R between the frequency of TdTOM^+^ DCs from (a) and (b) and frequency of all TdTOM^+^ TECs from Fig. 1b (*n*=8-10 mice from a minimum of 3 independent experiments). Statistical analysis in (c) and (d) was performed using a Pearson’s product-moment correlation, p≤0.05 = *, p≤0.01 = **, p≤0.001***, p<0.0001 = ****, ns = not significant.

It was previously described that the composition of TEC subsets differs with the age of mice. Specifically, the number of Aire^+^ mTEC^High^ cells has been shown to be decreased, while the number of Post-Aire mTECs gradually increased with age (Baran-Gale et al., 2020; Bornstein et al., 2018; Gray et al., 2006). Therefore, we compared the composition of the TEC subpopulations in TdTOM^+^ cells between young (4-6 weeks) and older (11-13 weeks) Csnβ^Cre^ mice to assess whether the changes in TdTOM composition in TECs would affect the frequency of TdTOM^+^ DCs. As expected, the population of TdTOM^+^ mTEC^High^ decreased, whereas TdTOM^+^ Post-Aire and Pre-post Aire mTECs increased with age (Figure 3b, left plot). Taking advantage of this phenomenon, we tested our prediction that the frequency of the TdTOM^+^ DC subsets would be altered in older Csnβ^Cre^ mice. Indeed, we observed a significant decrease in the frequency of cDC1s and Macs, along with an increase in XCR1^−^ aDC, cDC2, and pDC subsets (Figure 3b left graph). This data provides further evidence that CAT, as opposed to being mediated via random interactions, is a tightly regulated process that supports selective interactions between TEC and DC subsets.

To identify the predominant TEC-to-DC subsets trajectories of CAT, we performed a linear regression analysis of TdTOM^+^ TEC and TdTOM^+^ DC frequencies across Foxn1^Cre^, Defa6^iCre^, young Csnβ^Cre^, and older Csnβ^Cre^ mice (Figures 1e, 2g, and 3b). The data presented in Figure 3c confirmed the relatively narrow selectivity of each of the thymic DC subsets for certain TEC subset(s) from which they preferentially acquire antigens. Specifically, CAT to XCR1^+^ aDCs significantly correlated with the expression of TdTOM in mTECs^High^, Pre-post Aire, and Tuft mTECs, whereas XCR1^−^ aDCs aligned mostly with Pre-post Aire mTECs. cDC2s were the only subset that positively correlated with antigen production in Post-Aire mTECs. pDCs and Macs, and to a lesser extent cDC2s, correlated with mTEC^Low^. In addition, the Macs population significantly correlated with the expression of TdTOM in cTECs. This is consistent with the fact that the thymic Mac subset has been shown to preferentially reside in the thymic cortex (Breed et al., 2019). It is also important to emphasize that CAT to pDCs, Macs, and to lesser extend also to cDC2s, is highly affected by the frequency of total TdTOM^+^ TECs (Figure 3d). Thus, if the availability of TEC-derived antigens is limited, pDCs and Macs are outcompeted in CAT by other DC subsets. Surprisingly, the only positive correlation observed for cDC1 subset was with cTECs (Figure 3c). In this context, it was previously described that cDC1s predominantly acquired antigen from Aire^+^ mTEC^High^ subset (Lei et al., 2011; Perry et al., 2018). This discrepancy could be explained by the fact that cDC1s are the most represented population of TdTOM^+^ cells across all described models. Therefore, they show limited variability in their frequencies among TdTOM^+^ cells, which leaves little room for correlation in linear regression models. Therefore, we also performed a linear regression analysis of mTEC^High^ and cDC1 using only young and older Csnβ^Cre^ mice in which the variablility in the frequency of TdTOM^+^ cDC1 is higher (Figure 3b). This analysis indicated that cDC1s acquired antigen preferentially from mTEC^High^ cells (Supplementary Figure 4a). Remarkably, moDCs were the only DC subset that did not positively correlate with any of the TEC subsets (Figure 3c).

Together, this data confirms the hypothesis that CAT occurs between subsets of TECs and thymic DCs in a selective manner, with the exception of moDCs, which failed to reveal a preference for any subset of TECs.

### Thymic moDCs are the most efficient subset in repetitive CAT

Experiments that employed single-fluorescent protein transfer mouse models showed that most of the thymic CD11c^+^ subsets acquired antigens from more than one mTEC subset (Figure 3c). This poses the question of whether a single DC can take up antigens from several distinct TECs repetitively. To test this hypothesis, we utilized the Foxn1^Cre^Confetti^Brainbow2.1^ mouse model in which cytosolic RFP and YFP, and membrane CFP are expressed individually or in combination by TECs. The transfer of these fluorescent proteins to DCs (Figure 4a) was then measured. The expression of GFP, which should be present in the nucleus of Foxn1^Cre^Confetti^Brainbow2.1^ TECs (Snippert et al., 2010), was recently reported to be abrogated (Venables et al., 2019). By visualizing TECs from Foxn1^Cre^Confetti^Brainbow2.1^ and MHCII^eGFP^ mice, the latter used as a positive control, either separately or as a mixed cell suspension, confirmed that GFP is indeed absent in TECs from Foxn1^Cre^Confetti^Brainbow2.1^ mice (Supplementary Figure 5a). Given that YFP and RFP/CFP are expressed from mutually exclusive cassettes in Foxn1^Cre^Confetti^Brainbow2.1^ mice (Snippert et al., 2010), those TECs which express YFP do not express RFP and/or CFP and vice versa (Supplementary Figure 5b-d). Therefore, those DCs which were positive for both RFP and YFP must have obtained these antigens from two or more distinct TECs (Figure 4b). We found that this multi-antigen transfer occured nearly as frequently as the transfer from a single mTEC and that all CD11c^+^ APCs were involved in repetitive CAT. However, moDCs revealed the highest frequency of RFP^+^YFP^+^ cells which suggests a high level of promiscuity in targeting TEC subsets (Figure 4b and Supplementary Figure 5e).

**Figure 4.**
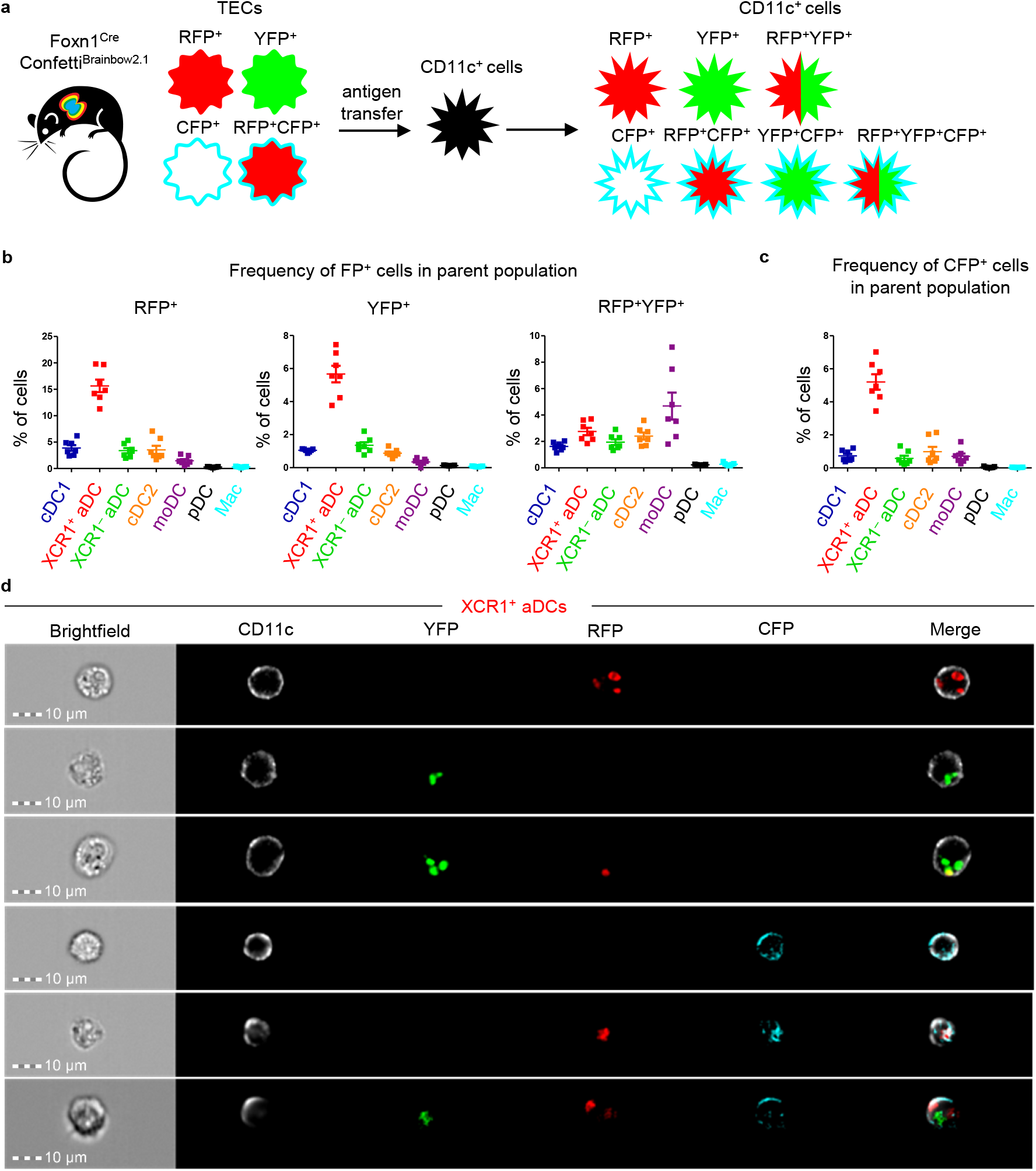
Thymic moDCs efficiently acquire antigens from two or more TEC cells in Foxn1^Cre^Confetti^Brainbow2.1^ mouse model. **(a)** Experimental design. **(b)** Quantification of the frequency of Fluorescent Protein^+^ (FP^+^) cells among the indicated DC subsets (mean ± SEM, *n*=7 mice from 3 independent experiments). **(c)** Quantification of the frequency of CFP^+^ cells among the indicated DC subsets (mean ± SEM, *n*=7 mice from 3 independent experiments). **(d)** Representative images from Imagestream analysis showing the localization of transferred FP in XCR1^+^ aDC from the thymus of Foxn1^Cre^Confetti^Brainbow2.1^ (n=2 independent experiments).

The Foxn1^Cre^Confetti^Brainbow2.1^ model also showed that the transfer of the CFP membrane antigen was observed less frequently than that of cytosolic antigens YFP and RFP. CFP transfer was largely mediated by XCR1^+^ aDCs which exhibited more than a 5-fold higher frequency of CFP positivity than any other CD11c^+^ subset (Figure 4c and Supplementary Figure 5f). Among the CFP^+^CD11c^+^ cell subsets, we also analyzed the co-acquisition of the other two fluorescent proteins (FPs) (Supplementary Figure 5g). As expected and consistent with their strong capacity to acquire FPs from more than one mTEC, the highest frequency of CFP^+^RFP^+^YFP^+^ cells were found in the moDC subset (Supplementary Figure 5g, right plot). There were only a few CFP^+^YFP^+^ cells observed in the CD11c^+^ subsets, which correlates with the overall low abundance of CFP single positive mTECs (Supplementary Figure 5b) and consequent low probability of a sequential encounter of YFP^+^ and CFP single positive TEC by CD11c^+^ cells. Since XCR1^+^ aDCs were, in general, the most potent DC subset in CAT in Foxn1^Cre^Confetti^Brainbow2.1^ mice, we imaged this subset with all possible FP^+^ variants using imaging flow cytometry (Figure 4d). It is of note, that CFP was in direct contrast to other FPs localized mainly to the plasma membranes of CAT-experienced XCR1^+^ aDCs.

Taken together, using Foxn1^Cre^Confetti^Brainbow2.1^ mice, we demonstrated that a single CD11c^+^ APC frequently acquired antigens from more than one mTEC and that the most potent subset in this repetitive CAT were moDCs. Moreover, we also showed that XCR1^+^ aDCs were very effective in the acquisition of both cytosolic and membrane-bound antigens.

### Thymic CD11c^+^ cells can share their antigens

Apart from the other CD11c^+^ APCs analyzed, the moDC subset showed no specific preference for any TEC subset in CAT (Figure 3c). This, together with their highest capacity among other CD11c^+^ subsets for repetitive CAT (Figure 4b) led us to test their possible involvement in the acquisition of antigens from other thymic CD11c^+^ cells. We performed a mixed BM chimera experiment in which irradiated CD45.1^+^CD45.2^+^ WT mice were reconstituted with a mix of BM (50:50) isolated from CD45.1^+^ WT and CD45.2^+^ CD11c^CRE^Rosa26^TdTOM^ mice (Figure 5a and Supplementary Figure 6a). Flow cytometric analysis showed that out of all CD45.1^+^CD11c^+^ cells, approximatelly 0,75% acquired TdTOM from CD45.2^+^CD11c^+^ cells (Figure 5b-c). While the contribution of both aDC subsets and cDC2s to CAT was robust, the highest frequency of TdTOM^+^ cells was found among the moDC subset (Figure 5d and Supplementary Figure 6b). Thus, thymic CD11c^+^ cells, especially moDCs, acquire antigens not only from TECs but from other CD11c^+^ cells as well.

**Figure 5.**
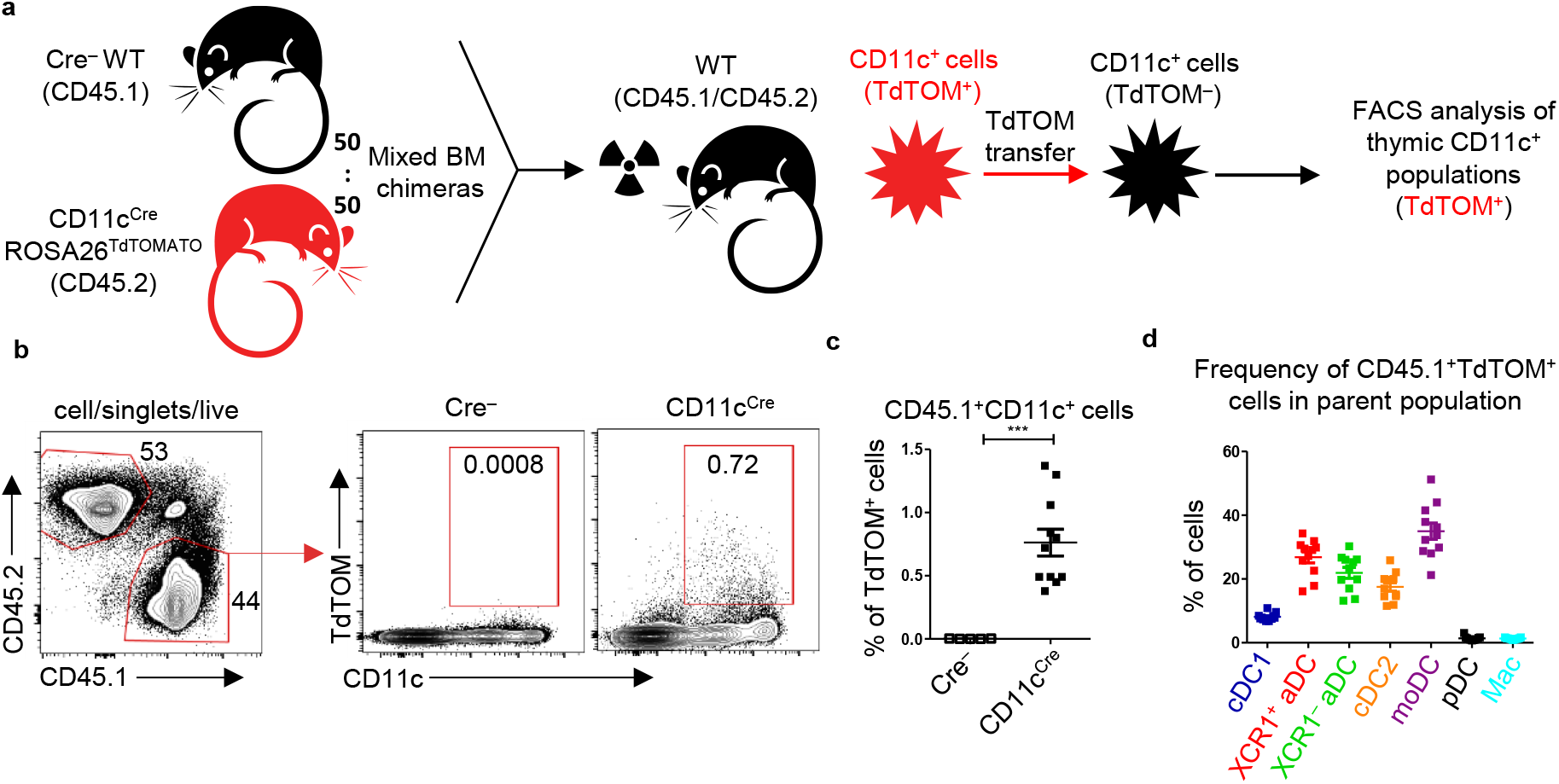
Thymic CD11c+ cells can share their antigens between each other. **(a)** Experimental design. **(b)** Representative flow cytometry plots showing the frequency of CD45.1^+^CD11c^+^TdTOM^+^ cells among MACS-enriched CD11c^+^ thymic cells from mixed, bone marrow chimeras (50:50) of WT (CD45.1^+^) and CD11c^Cre^ROSA26^TdTOM^ (CD45.2^+^) mice. **(c)** Quantification of CD45.1^+^TdTOM^+^CD11c^+^ cells from (b) (mean ± SEM, *n*=11 mice from 2 independent experiments). Statistical analysis was performed using unpaired, two-tailed Student’s t-test, p≤0.001***. **(d)** Quantification of the frequency of TdTOM^+^ cells among the indicated DC subsets from reconstituted mice described in (a) (mean ± SEM, n=11 mice from 2 independent experiments).

Together, this data demonstrates that the acquisition of antigens by the thymic population of CD11c^+^ cells is not restricted to TEC subsets but is extended to other cell-subtypes, mainly to their own CD11c^+^ cells. Remarkably, among all thymic DCs, moDCs were the most efficient in this special type of “cannibalistic” CAT.

## Discussion

This study, which has been based on initial observations by others (Lancaster et al., 2019; Mouri et al., 2017; Perry et al., 2018), confirmed that CAT, i.e. TEC-to-DC antigen-spreading, is not a random process. Using these studies along with reports concerning the heterogeneity of thymic APCs as a foundation, we have provided detailed insight into how particular subsets of TECs and thymic APC are interconnected in the transfer of TEC-produced antigens. Specifically, utilizing several murine genetic models which allowed the tracking of TEC-produced antigen, we determined that CAT is mediated predominantly by preferential pairing between the following TECs and CD11c^+^ DC subsets: (i) mTEC^Low^ to pDC and Mac, (ii) mTEC^High^ to cDC1 and XCR1^+^ aDC, (iii) Pre-post Aire mTEC to XCR1^+^ and XCR1^−^ aDC, (iv) Post-Aire mTEC to cDC2, and (v) Tuft mTEC to XCR1^+^ aDC. These CAT trajectories, which are depicted in Figure 6a, argue in favor of a model of preferential pairing in thymic antigen transfer. However the antigen acquisition by pDCs and Macs is effective only when the antigen is abundant. In addition, we also report that thymic moDCs, which do not exhibit subset specificity in CAT, generally obtain antigen from multiple cellular sources of thymic TECs as well as CD11c^+^ DC subsets.

**Figure 6.**
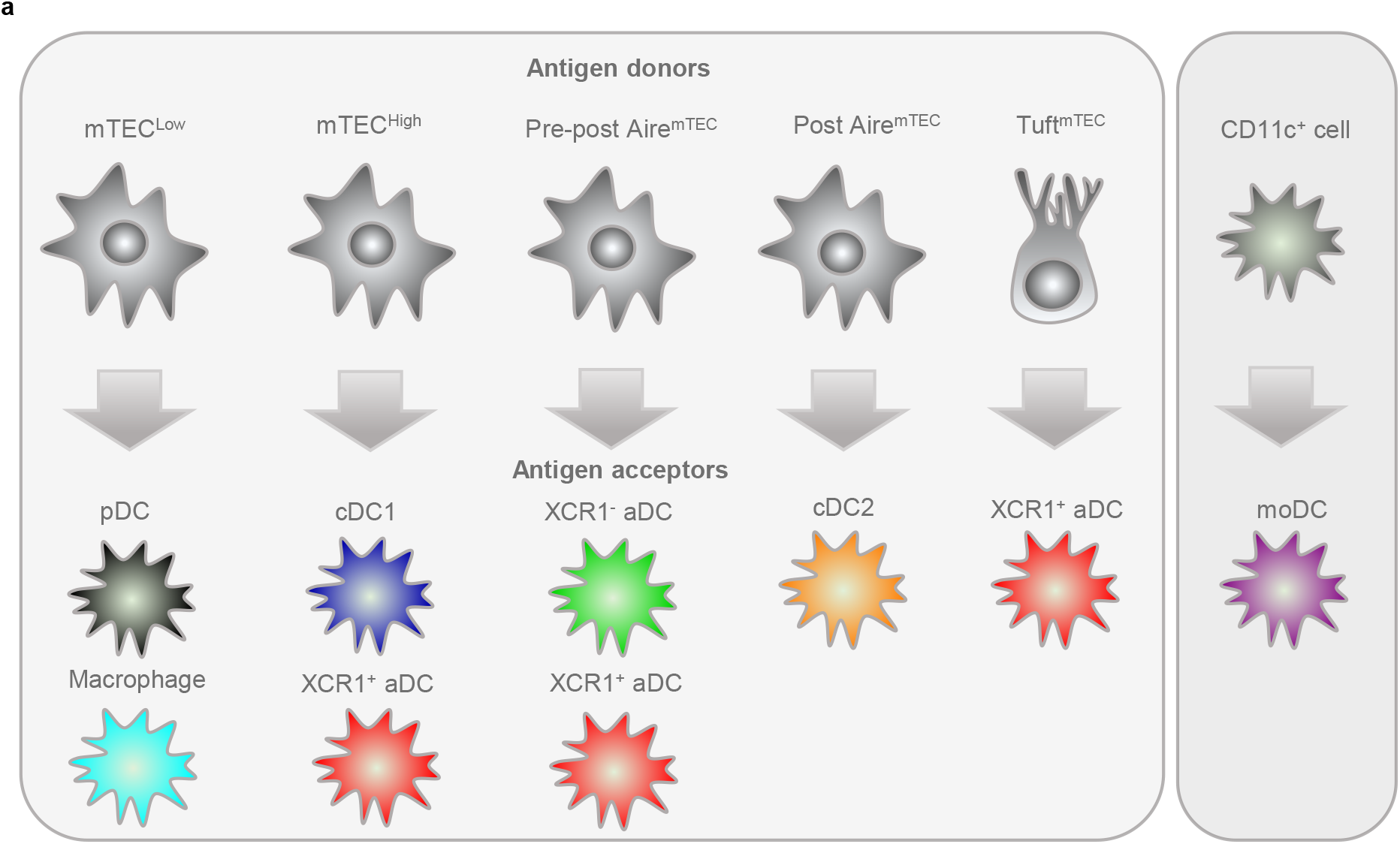
Proposed model of preferential pairing in CAT. **(a)** Based on the data presented in this study, we postulate that phenotypically defined subsets of thymic CD11c^+^ cells preferentially and predictably acquire antigens from distinct subsets of developmentally-related TECs (left panel). Our data also suggests that thymic moDC do not specifically prefer any particular subset of TECs and simultaneously are efficient in acquiring antigens from other CD11c^+^ APCs (right panel). This suggests that moDCs generally act as cells that scavenge apoptotic TECs and APCs in the thymic medulla.

In this study, we confirmed a high level of internal TEC heterogeneity which could be divided into a minimum of six distinct subsets (Baran-Gale et al., 2020; Bautista et al., 2021; Bornstein et al., 2018; Dhalla et al., 2020; Wells et al., 2020). Since the majority of these subsets are developmentally related to each other (Bornstein et al., 2018; Metzger et al., 2013; Miller et al., 2018), our Cre-based ROSA26^TdTOM^ mouse models (Figure 1a) can be employed as lineage tracing systems for tracking developmental relationships between mTEC subsets. It has been reported that TdTOM expression in the Csnβ^Cre^ROSA26^TdTOM^ mouse model is detected in a small proportion of mTEC^Low^, in most mTEC^High^, Post-Aire mTEC, and Tuft mTEC subsets (Bornstein et al., 2018). This is consistent with our data (Figure 1d), which suggests that Csnβ is expressed by a specific population of mTEC^Low^ progenitors that further differentiate into mTECs^High^ cells and later into their progeny. In contrast, the TdTOM expression in Defa6^iCre^ROSA26^TdTOM^ should be specifically attributed to Aire^+^ mTEC^High^ subset and their Post-Aire progeny, since the expression of defensins in the thymus is highly dependent on Aire (Filipp et al., 2018). Despite that, we see the TdTOM expression also in the small LY6D^−^ population of mTEC^Low^ (Figure 1d). Since several distinct subpopulations of Post-Aire mTEC were detected (Dhalla et al., 2020), we hypothesized that Cre recombination in mTEC^Low^ reflects the presence of LY6D^−^ population of Post-Aire cells than Defa6 locus activation in Aire^−^ mTEC^Low^ progenitors. Thus, the significant correlation in CAT between mTEC^Low^ and cDC2 subsets could be influenced by this phenomenon, since cDC2s were shown to acquire the antigen mostly from Post-Aire mTECs (Figure 3c). It is also important to emphasize that TdTOM^+^ Tuft mTECs were enriched in the Defa6^iCre^ mouse model compared to Csnβ^Cre^ (Figure 1e). This suggests that thymic Tuft cells are descendants of Aire^+^ mTEC^High^ subset (Miller et al., 2018).

The development of novel gating strategies has allowed us to reveal the substantial heterogeneity of thymic DCs which could be divided into phenotypically and functionally distinct subsets (Li et al., 2009; Park et al., 2020; Vobořil et al., 2020). Our data points to at least seven subtypes of CD11c^+^ cells that are capable of antigen acquisition from different subsets of TECs. i.e. cDC1, XCR1^+^ aDC, XCR1^−^ aDC, cDC2, moDC, pDC, and a population of Macs (Figure 2e and Supplementary Figure 3a-b). Among them, we have phenotypically defined two novel subsets of thymic aDCs, which are marked by the overexpression of the chemokine receptor, CCR7. Notably, it has been reported that the expression of CCR7 defines the population of XCR1^+^CCR7^+^ cDC1s which are considered to be the progeny of XCR1^+^CCR7^−^ cDC1s (Ardouin et al., 2016). However, since these CCR7^+^ cDC1s express several molecules that are not only associated with the cDC1 signature, such as *Batf3, Cd8α, Ly75*, or *Cadm1* (Vobořil et al., 2020) but also molecules that have been attributed to the population of aDCs (*Il12b, Il15, Il15ra Cd274, Cd70, Cd40, Tnfrsf4*) (Ardouin et al., 2016; Park et al., 2020) we defined and renamed this subset as XCR1^+^ aDC. Remarkably, these cells are the most efficient DC subset in CAT, even when compared to cDC1s (Supplementary Figure 2b, Figure 4b) (Ardouin et al., 2016). It was recently suggested that the differentiation of XCR1^+^ aDCs from cDC1s is driven by the uptake of apoptotic cells (Maier et al., 2020). Since CAT has been shown to be mediated mostly by the endocytosis of apoptotic bodies (Koble & Kyewski, 2009; Perry et al., 2018), the differentiation of XCR1^+^ aDCs in the thymus is consistent with being driven by CAT. Thus, the grounds for the correlation between mTEC^High^ and XCR1^+^ aDCs in TdTOM antigen transfer could be found in the fact that mTEC^High^ transfer antigen to XCR1^+^ cDC1 which further differentiate into XCR1^+^ aDC cells (Ardouin et al., 2016; Maier et al., 2020). In this context, it is also important to emphasize that the transcriptional signature of XCR1^−^ aDCs is more similar to cDC2 (e.g. *Sirpa* and *Pdcd1lg2*) than cDC1 subset (Park et al., 2020). By the same token, this suggests that antigen transfer into cDC2s induces their differentiation into XCR1^−^ aDCs.

Using linear regression analysis of TdTOM^+^ TECs and DCs frequencies from all three mouse models, we identified two subsets of CD11c^+^ cells, cDC1 and moDC, that exhibited limited or no correlation with TEC subsets in TdTOM transfer. cDC1s were observed to correlate with cTECs (Figure 3c). This is contradictory to previously published data which described the antigen uptake by cDC1s specifically from Aire^+^ mTEC^High^ (Lei et al., 2011; Perry et al., 2018). The data shown in Supplementary Figure 4a supports this conclusion. As briefly stated in the results section, we view the correlation in CAT between cDC1 and cTEC subsets as an artifact of the linear regression model because of the variability in TdTOM^+^ frequencies of these two subsets across all Cre-based ROSA26^TdTOM^ models remained, for the most part, unchanged (Figure 3a). We also based this conclusion on the fact that cDC1s are preferentially localized to the thymic medulla, whereas cTECs take up residence in the thymic cortex, a condition which is not conducive for cell interaction (Klein et al., 2014). Additionally, since CAT has been shown to be cell contact-dependent (Kroger et al., 2017; Perry et al., 2018) and XCL1-XCR1 chemotactic axis is essential for CAT between Aire^+^ mTEC^High^ and cDC1 subsets (Lei et al., 2011), we favor the scenario that cDC1s acquire antigen preferentially from mTEC^High^ subset and not cTECs (Figure 6a).

The second subset of thymic CD11c^+^ cells, which failed to show a correlation with any TEC-subset in CAT consisted of the moDCs. Interestingly, while moDCs are very potent in CAT (Figure 2e and Supplementary Figure 2b), their capacity can be further enhanced under inflammatory conditions (Vobořil et al., 2020). We demonstrated that among other thymic DCs, moDC subset were the most efficient in repetitive CAT (Figure 4b and Supplementary Figure 5g). This, along with their ability to effeciently acquire antigen from other CD11c^+^ APCs (Figure 5d), is a testament to their important function in central tolerance (Park et al., 2020; Vobořil et al., 2020). Since thymic moDCs were shown to express a plethora of different chemokines and scavenger receptors (Park et al., 2020; Vobořil et al., 2020), we propose that these characteristics correlate with their high competence in regulated migration and phagocytic activity compared to other DC subsets (Croxford et al., 2015).

In conclusion, using novel gating strategies for the identification of multiple TEC subsets which produce TdTOM antigen and tracking of its transfer into phenotypically defined thymic CD11c^+^ APC subsets has allowed us to define preferential antigen trajectories which mediate CAT. Our data shows that XCR1^+^ aDCs are the most potent subset in the acquisition of TEC-derived antigens. It also characterizes the moDC subset as the most efficient in the acquisition of antigen from multiple TECs as well as DCs. Taken together, our work proposes that CAT relies on a cellular interaction network with preferential partnerships between defined subtypes of TECs and DCs. This, in turn, suggests that the indirect presentation of antigens from developmentally related but phenotypically and functionally distinct types of TECs is ascribed to different subsets of thymic DCs. However, how these cell-to-cell preferential interactions which are the underlying characteristics of CAT facilitate the processes of central tolerance, such as the deletion of self-reactive clones of T-cells or their conversion to Tregs awaits its resolution. Although this study suggests that CAT is a deterministic process, the molecules and mechanisms that determine TEC-to-DC cell-cell interactions remain to be identified.

## Materials and Methods

### Mice

All mouse models used in this study were of C57BL/6J genetic background and housed under SPF conditions at the animal facility of the Institute of Molecular Genetics (IMG) in Prague. All animal experiments were approved by the ethical committee of the IMG and the Czech Academy of Sciences. C57BL/6J, Foxn1^Cre^ (B6(Cg)-Foxn1^tm3(cre)Nrm^/J) (Gordon et al., 2007), Ly5.1 (B6.SJL-Ptprc^a^ Pepc^b^/BoyJ) (Janowska-Wieczorek et al., 2001), and CD11c^Cre^ (B6.Cg-Tg(Itgax-cre)1-1Reiz/J) (Caton et al., 2007) mice were purchased from Jackson Laboratories. Csnβ^Cre^ mice (Bornstein et al., 2018) were kindly provided by J. Abramson (Department of Immunology, Weizmann Institute of Science, Rehovot, Israel). Defa6^iCre^ mice (Adolph et al., 2013) were kindly provided by R. S. Blumberg (Division of Gastroenterology, Department of Medicine, Brigham and Women’s Hospital, Harvard Medical School, Boston, Massachusetts). Rosa26^TdTOMATO^ mice (B6;129S6-Gt(ROSA)26Sor^tm14(CAG-tdTomato)Hze^/J) (Madisen et al., 2010) were provided by V. Kořínek (IMG, Prague, Czech Republic). Confetti^Brainbow2.1^ (Gt(ROSA)26Sor^tm1(CAG-Brainbow2.1)Cle^/J) (Snippert et al., 2010) mice were provided by the Czech Center for Phenogenomics (IMG, Vestec, Czech Republic). MHCII^eGFP^ mice (Boes et al., 2002) were provided by J. Černý (Department of Cell Biology, Faculty of Science, Charles University, Prague). All mice were fed an Altromin 1314 IRR diet. Reverse osmosis filtered and chlorinated water was available to the animals ad libitum. All mice were bred in an environment in which the temperature and humidity of 22 ± 1°C and 55 ± 5%, respectively was constant and under a 12 h oscillating light/dark cycle. Prior to tissue isolation, mice were euthanized by cervical dislocation.

### Tissue preparation and cell isolation

Thymic tissue was extracted using forceps, cut into small pieces, and enzymatically digested with 0.1 mg*ml^− 1^ Dispase II (Gibco) dissolved in RPMI. Pieces of thymic tissue were pipetted up and down several times using a pipet tip that had been cut and incubated in a shaker at 800 rpm for 10 min at 37°C. This procedure was repeated ∼5 times to completely dissolve the tissue. The supernatant was collected and the enzymatic reaction was stopped by adding 3% FCS and 2 mM EDTA. To isolate thymic epithelial cells (TECs), isolated cells were MACS-depleted of CD45^+^ cells using CD45 microbeads (Miltenyi). After depletion, the suspension was spun down (4 °C, 300 g, 10 min) and the resulting pellet was resuspended in ACK lysis buffer for 2 min to deplete erythrocytes. To isolate thymic DCs and macrophages, MACS enrichment for CD11c^+^ cells was performed using CD11c biotin-conjugated antibody (eBioscience) and Ultrapure Anti-Biotin microbeads (Miltenyi).

### Flow cytometry analysis and cell sorting

Cell staining for flow cytometry (FACS) analysis and sorting was performed at 4 °C, in the dark, for 20–30 min, with the exception of anti-CCR7 antibody (Biolegend) staining which required incubation at 37°C for a minimum of 30 min. To exclude dead cells, either Hoechst 33258 (Sigma-Aldrich) or viability dye eFluor 506 (eBioscience) was used. FACS analysis of TECs and DCs was performed using BD™ LSR II and BD™ FACSymphony A5 cytometers, respectively. A BD™ FACSAria IIu sorter was used for cell sorting. BD FACSDiva™ Software and FlowJO V10 software (Treestar) were used for FACS data analysis. For the purpose of tSNE analysis, the same amount of CD11c^+^ TdTOM^+^ cells from each model was concatenated by using the FlowJO concatenate function. The final tSNE was calculated by FlowJO opt–SNE plugin. The entire list of FACS staining reagents is provided in Supplementary Table 1.

### Imaging flow cytometry

Imaging flow cytometry (Imagestream) was performed using AMNIS ImageStream X MkII at the Center for Preclinical Imaging (CAPI) in Prague. Imaged XCR1^+^ aDC were isolated from Foxn1^Cre^Confetti^Brainbow2.1^ mice, stained for their CD11c, XCR1, and CCR7 markers, and sorted as RFP^+^ and/or YFP^+^ and/or CFP^+^. The data was acquired via Imagestream with 40x magnification. Ideas 6.1 software (AMNIS) was used for data analysis.

### Confocal and spinning disk microscopy

To test GFP expression in TECs from Foxn1^Cre^Confetti^Brainbow2.1^ mice (Supplementary Fig. 5a), thymic cells from Foxn1^Cre^Confetti^Brainbow2.1^ and MHCII^eGFP^ mice were MACS-depleted of CD45^+^ fraction and imaged on a Leica TCS SP5 AOBS Tandem confocal microscope using the HCX PL APO 10x/0.40 DRY CS; FWD 2.2; CG 0.17 | BF, POL objective. To visualize TEC fluorescent variants from Foxn1^Cre^Confetti^Brainbow2.1^ mice (Supplementary Fig. 5d), CD45^+^EpCAM^+^ TECs were sorted as RFP^+^ and/or YFP^+^ and/or CFP^+^ and visualized with a Andor Dragonfly 503 spinning disk confocal microscope using HCX PL APO 63x/1.40-0.6 OIL λB; FWD 0.12; CG 0.17 | BF, POL, DIC objective.

### Bone marrow chimeras

Bone marrow was flushed out from the femur and tibia of Ly5.1 (CD45.1^+^; Fig. 5, Supplementary Fig. 2d, e, f and 6) or CD11c^CRE^Rosa26^TdTOMATO^ (CD45.2^+^; Fig. 5 and Supplementary Fig. 6) mice using a syringe with 26g needle. Isolated cells were depleted of erythrocytes with ACK lysis buffer. Recipient mice were sublethally irradiated with 6 Gy and reconstituted with 2 × 10^6^ Ly5.1 BM cells in the case of Foxn1^Cre^/Csnβ^Cre^/Defa6^iCre^ROSA26^TdTOMATO^ mice (Supplementary Fig. 2d, e, f) or with 2 × 10^6^, 50:50 mixed Ly5.1:CD11c^CRE^Rosa26^TdTOMATO^ BM cells in the case of C57BL/6J Ly5.1 mice (CD45.1^+^CD45.2^+^; Fig. 5 and Supplementary Fig. 6). Three weeks after irradiation, the BM reconstitution was verified by the staining of blood with anti-CD45.1 and CD45.2 antibodies. Mice were subjected to further analysis 6 weeks after irradiation if the BM reconstitution exceeded 80% (Supplementary Fig. 2d, e, f) or was between 40–60% within both CD45.1^+^ and CD45.2^+^ cell compartments (Fig. 5 and Supplementary Fig. 6).

### Statistical analysis

Statistical analysis and graph layouts were performed using Prism 5.04 software (GraphPad). Linear regressions were calculated using R 3.6.2. (R core team 2019). The statistical tests used for data analysis are indicated in figure legends.

## Acknowledgments

We would like to thank the members of the flow cytometry facility, Z. Cimburek, and M. Šíma, from IMG in Prague for technical support and I. Novotný for assistance with microscopic experiments. We also thank M. Báječný for technical assistance with Imaging flow cytometry. We are indebted to J. Abramson for providing the Csnβ^Cre^ mice, V. Kořínek for ROSA26^TdTOMATO^ mice, and the Czech Center for Phenogenomics for Confetti^Brainbow2.1^ mice. This work was supported by Grant 20-30350S from GACR. J.B. was supported by Grant 836119 from the Charles University Grant Agency (GA UK). J.D. was supported by a PRIMUS grant (Primus/21/MED/003) from Charles University and a Czech Science Foundation JUNIOR STAR grant (GAČR 21-22435M). T.B. was partially supported by Grant RVO: 68378050-KAV-NPUI.

## Author contributions

M.V. and J.B. co-designed and conducted the majority of the experiments and wrote the manuscript. T.B. performed some of the experiments and provided technical and intellectual help. J.D. conducted Imaging flow cytometry experiment and helped with the preparation of the manuscript. O.B. performed microscopic experiments and M.D. and J.M. provided technical support for the work and manuscript editing. R.B. provided Defa6^iCre^ mice. D.F. co-designed experiments, supervised research and edited the manuscript.

## Supplementary information

**Supplementary Figure 1.**
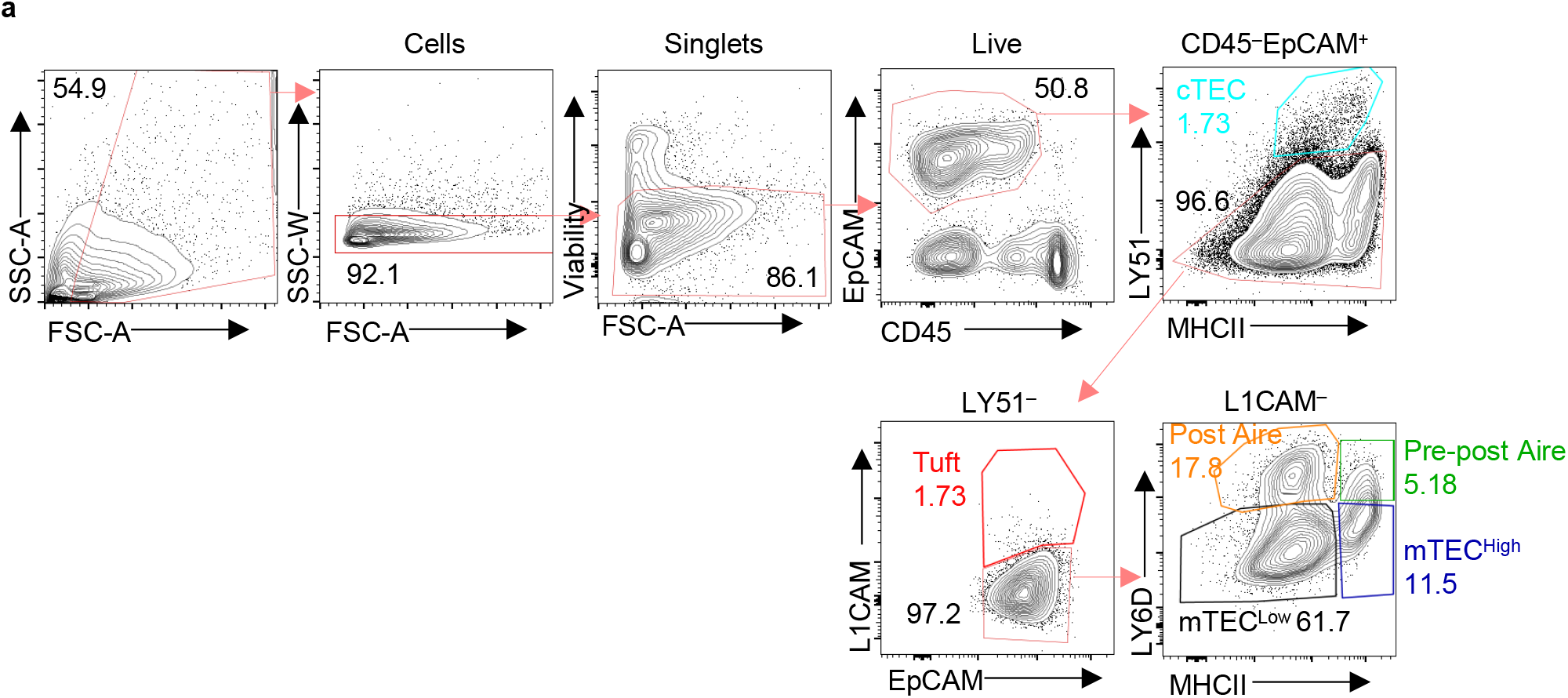
Related to Figure 1. Gating strategy of thymic epithelial cell populations. **(a)** Complete gating strategy for the distiction of TEC populations. The thymic cell fraction was MACS enriched for CD45^−^ cells and sequentially gated as singlets, live, and CD45^−^ EpCAM^+^ cells. The fraction of isolated TECs was then gated as cTECs (LY51^+^), Tuft mTECs (LY51^−^L1CAM^+^), and mTECs (LY51^−^L1CAM^−^). mTECs consists of four major populations: mTEC^Low^ (MHCII^Low^LY6D^−^), mTEC^High^ (MHCII^High^LY6D^−^), Pre-post Aire mTECs (MHCII^High^LY6D^+^), and Post Aire mTECs (MHCII^High^Ly6D^High^).

**Supplementary Figure 2.**
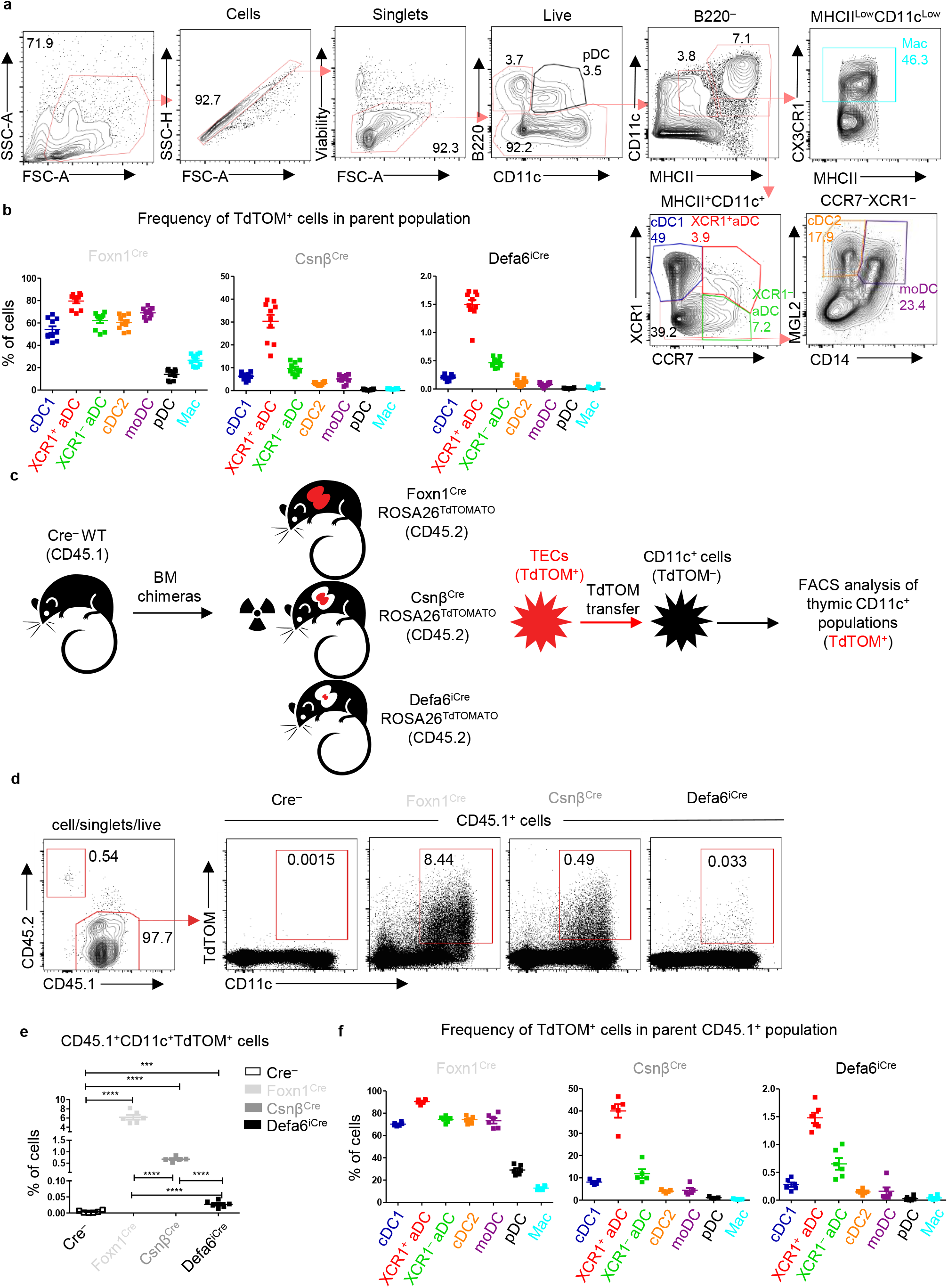
Related to Figure 2. Antigen transfer of TdTOMATO to thymic dendritic cells. **(a)** Complete gating strategy for the isolation of thymic DC populations. The thymic cell fraction was MACS enriched for CD11c^+^ cells and sequentially gated as singlets and live cells. This cell fraction was then depleted of pDCs (B220^+^CD11c^Low^) and divided into CD11c^+^MHCII^+^ and CD11c^Low^MHCII^Low^ populations. CD11c^+^MHCII^+^ cells represent the major thymic DC populations: cDC1 (XCR1^+^CCR7^−^), XCR1^+^ aDC (XCR1^+^CCR7^+^), XCR1^−^ aDC (XCR1^−^CCR7^+^), cDC2 (XCR1^−^CCR7^−^MGL2^+^CD14^−^), and moDC (XCR1^−^CCR7^−^ MGL2^+^CD14^+^). CD11c^Low^MHCII^Low^ cells contain CX3CR1^+^, a macrophage-like population (Mac). Historically, SIRPα gating was used to distinguish cDC1 from cDC2 subsets. Since the XCR1 and CCR7 gating enables us to distinguish several subsets of thymic DCs, we omitted SIRPα from our gating strategy. (**b)** Quantification of the frequencies of TdTOM^+^ cells among the indicated DC subsets (mean ± SEM, *n*=10 from minimum of 3 independent experiments). **(c)** Experimental design. **(d)** Representative flow cytometry plots compairing the frequency of CD45.1^+^TdTOM^+^CD11c^+^ cells among MACS-enriched CD11c^+^ thymic cells from the mouse models described in (c). **(e)** Quantification of CD45.1^+^TdTOM^+^CD11c^+^ cells from (d) (mean ± SEM, *n*=5-6 mice from 2 independent experiments). **(f)** Quantification of the frequency of TdTOM^+^ cells among the indicated DC subsets from mice described in (c) (mean ± SEM, n= 5-6 mice from 2 independent experiments).

**Supplementary Figure 3.**
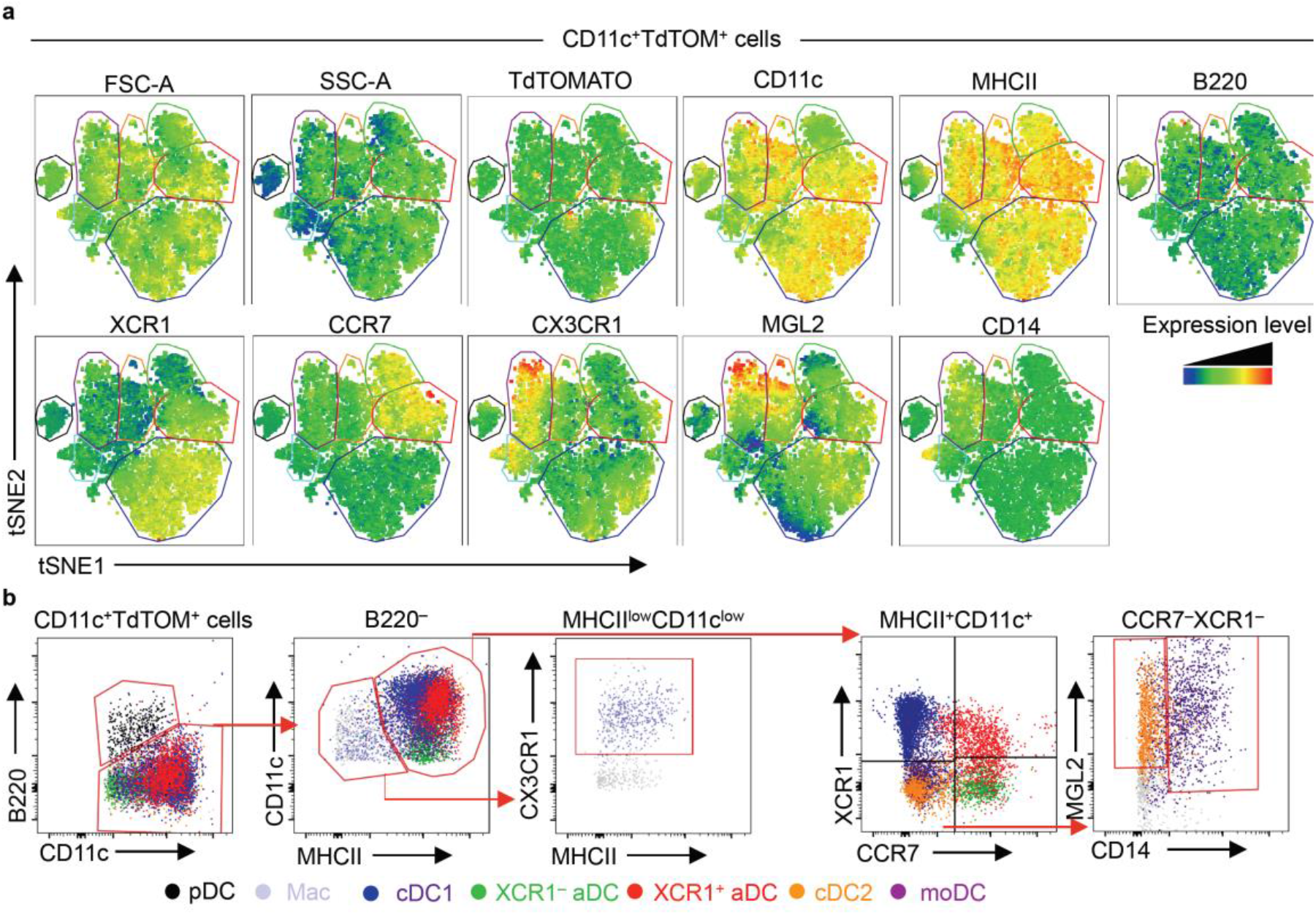
Related to Figure 2. Thymic dendritic cell gating strategy defined by flow cytometry tSNE analysis. **(a)** Heat map generated from flow cytometry tSNE analysis of TdTOM^+^CD11c^+^ cell populations from Figure 2d. tSNE analysis was performed using FlowJO software, based on the FSC-A, SSC-A, TdTOMATO, CD11c, MHCII, B220, XCR1, CCR7, CX3CR1, MGL2, and CD14. **(b)** Back-gating of TdTOM^+^CD11c^+^ populations defined in (a), onto the CD11c^+^ gating strategy described in Supplementary Figure 2a.

**Supplementary Figure 4.**
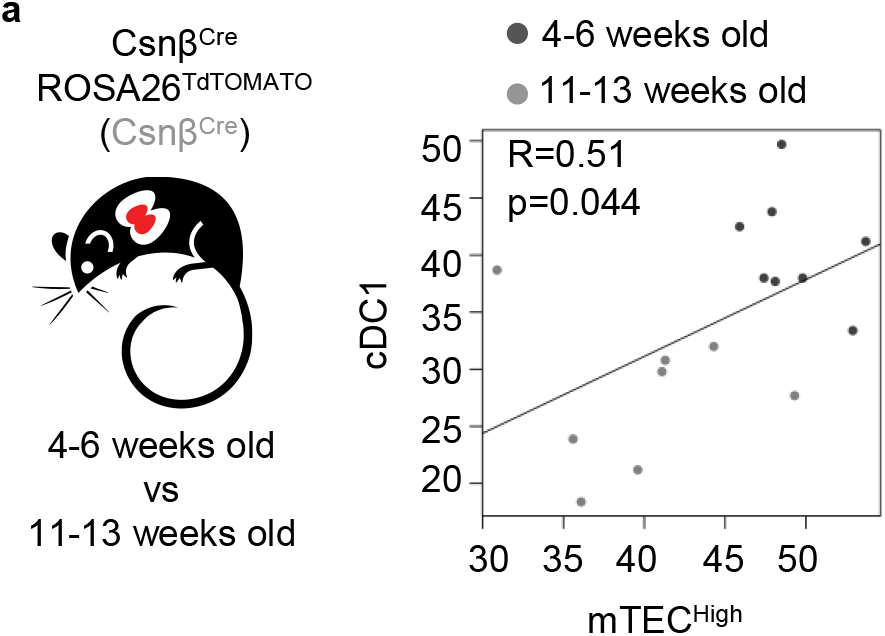
Related to Figure 3. The TdTOMATO antigen transfer to cDC1 subset correlates with its expression in mTEC^High^. **(a)** Linear regression (R) between frequencies of TdTOM^+^ mTEC^High^ and TdTOM^+^ cDC1 from Csnβ^Cre^ mice described in Figure 3b (*n*=8 mice from minimum of 3 independent experiments). Statistical analysis was performed by Pearson’s product-moment correlation.

**Supplementary Figure 5.**
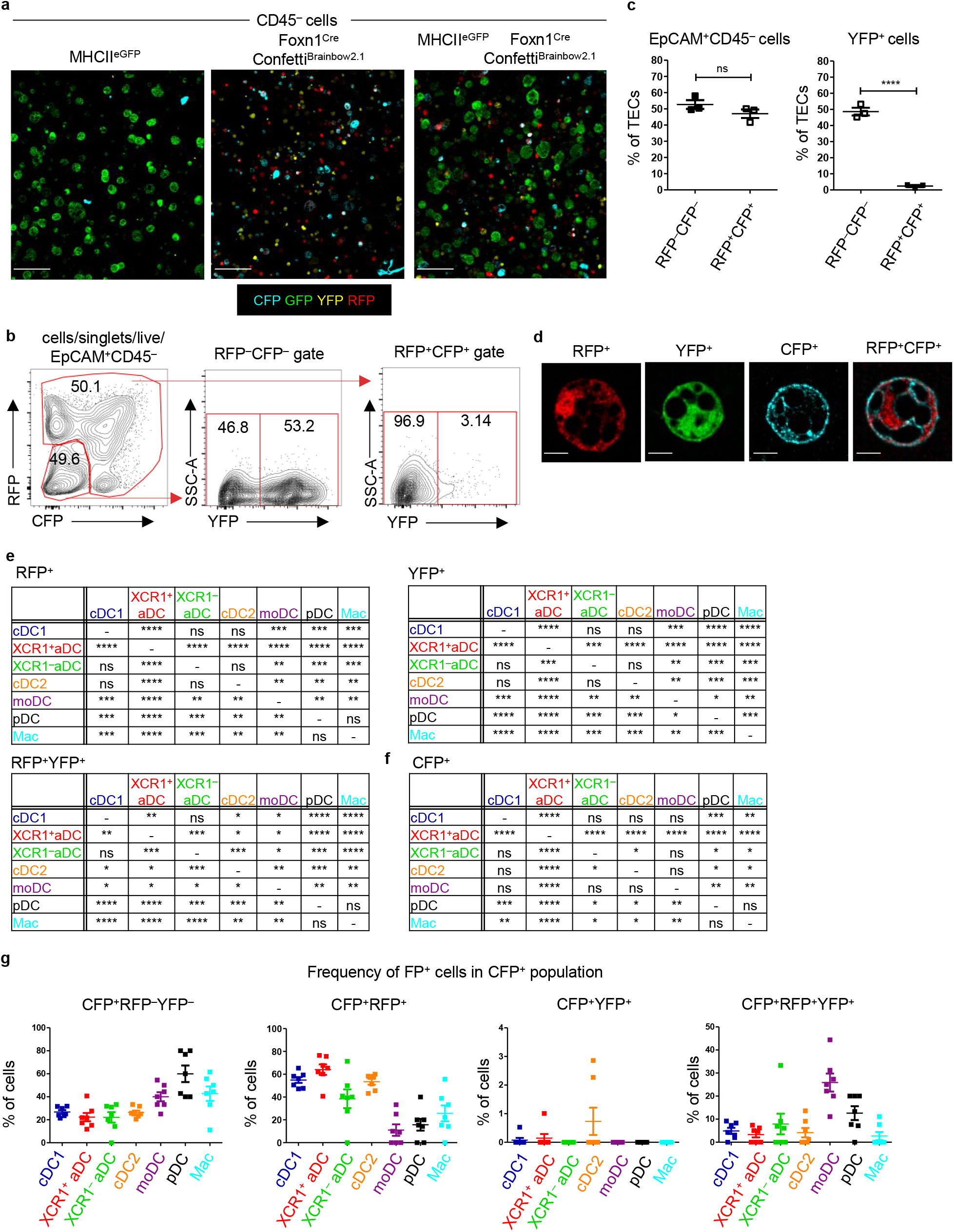
Related to Figure 4. Foxn1^Cre^Confetti^Brainbow2.1^ as a model of thymic cooperative antigen transfer. **(a)** Representative microscopic images of sorted TECs from MHCII^eGFP^ (left panel), Foxn1^Cre^Confetti^Brainbow2.1^ (middle panel) and a mixed population of TECs isolated from both MHCII^eGFP^ and Foxn1^Cre^Confetti^Brainbow2.1^ (right panels) mouse models. **(b)** Representative flow cytomentry plots showing the frequency of YFP, RFP and CFP^+^ CD45^−^ EpCAM^+^ TECs. **(c)** Quantification of FP^+^ cells from (b) (mean ± SEM, *n*=3 mice from 2 independent experiments). **(d)** Representative microscopic images of all TEC variants from the model described in Figure 5a. **(e), (f)** Statistical analysis of the frequency of FP^+^ cells among the indicated DC subsets from Figure 4b and c (*n*=7 mice from 3 independent experiments). Analysis was performed using a paired, two-tailed Student’s t-test, p≤0.05 = *, p≤0.01 = **, p≤0.001***, p<0.0001 = ****, ns = not significant. **(g)** Quantification of the frequency of FP^+^ cells among the CFP^+^ DC subsets (mean ± SEM, *n*=7 from 3 independent experiments).

**Supplementary Figure 6.**
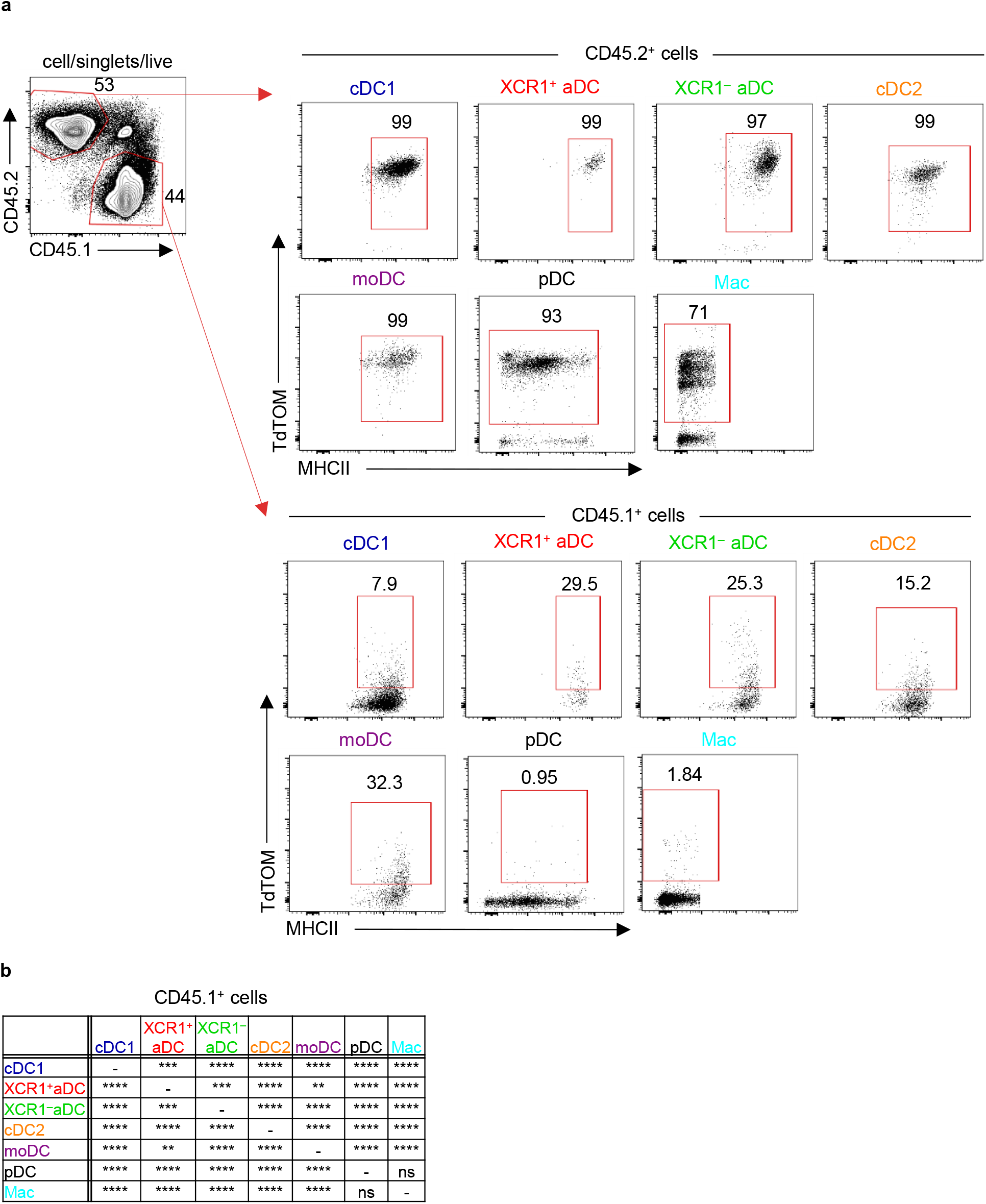
Related to Figure 5. Thymic CD11c^+^ cells can share their antigens between each other. **(a)** Representative flow cytometry plots showing gating and frequency of CD11c^+^TdTOM^+^ cells from mixed, bone marrow chimera of WT (CD45.1^+^, lower panel) and CD11c^Cre^ROSA26^TdTOMATO^ (CD45.2^+^, upper panel) mice. **(b)** Statistical analysis of the frequency of TdTOM^+^ cells among the indicated DC subsets from Figure 5d (*n*=11 mice from 2 independent experiments). Analysis was performed using a paired, two-tailed Student’s t-test, p≤0.01 = **, p≤0.001***, p<0.0001 = ****, ns = not significant.

**Supplementary Table 1.** List of antibodies.

| **Antibody** | **Manufacturer** | **Clone** | **Catalogue number** | **Dilution** |
| --- | --- | --- | --- | --- |
| Anti-mouse CCR7-APC | BioLegend | 4B12 | cat# 120108 | 1:100 |
| Anti-mouse CCR7-PE/Cy7 | BioLegend | 4B12 | cat# 120124 | 1:200 |
| Anti-mouse CD11c-APC/Cy7 | BioLegend | N418 | cat# 117324 | 1:200 |
| Anti-mouse CD11c-Biotin | eBioscience | N418 | cat# 13-0114-82 | 1:100 |
| Anti-mouse CD14-APC | BioLegend | Sa2-8 | cat# 123312 | 1:100 |
| Anti-mouse CD14-FITC | BioLegend | Sa2-8 | cat# 123308 | 1:100 |
| Anti-mouse CD301b (Mgl2)-PE/Cy7 | BioLegend | URA-1 | cat# 146807 | 1:200 |
| Anti-mouse CD326 (EpCAM)-PE/Cy7 | BioLegend | G8.8 | cat# 118215 | 1:3000 |
| Anti-mouse CD45.1-PerCP/Cy5.5 | BioLegend | A20 | cat# 110727 | 1:150 |
| Anti-mouse CD45.2-FITC | BioLegend | 104 | cat# 109806 | 1:150 |
| Anti-mouse CD45-BV605 | BioLegend | 30-F11 | cat# 103155 | 1:100 |
| Anti-mouse CX3CR1-BV421 | BioLegend | SA011F11 | cat# 149023 | 1:200 |
| Anti-mouse I-A/I-E-BV711 | BioLegend | M5/114.15.2 | cat# 107625 | 1:500 |
| Anti-mouse I-A/I-E-PB | BioLegend | M5/114.15.2 | cat# 138606 | 1:500 |
| Anti-mouse L1CAM-APC/Cy7 | Novus Biologicals | UJ127.11 | cat# 2682APCCY7 | 1:50 |
| Anti-mouse Ly-51-AF647 | BioLegend | 6C3 | cat# 108311 | 1:200 |
| Anti-mouse Ly6D-FITC | BioLegend | 49-H4 | cat# 107620 | 1:200 |
| Anti-mouse/human CD45R/B220-BV785 | BioLegend | RA3-6B2 | cat# 103246 | 1:100 |
| Anti-mouse/rat XCR1-APC | BioLegend | ZET | cat# 148206 | 1:200 |
| Anti-mouse/rat XCR1-BV421 | BioLegend | ZET | cat# 148216 | 1:200 |
| Anti-mouse/rat XCR1-PerCP/Cy5.5 | BioLegend | ZET | cat# 148208 | 1:200 |
| Fixable Viability Dye-eFluor 506 | eBioscience | - | cat# 65-0866-14 | 1:300 |
| Streptavidin-APC/Cy7 | BioLegend | - | cat# 405208 | 1:500 |

